# Detecting substrate glycans of fucosyltransferases on glycoproteins with fluorescent fucose

**DOI:** 10.1101/2020.01.28.919860

**Authors:** Zhengliang L Wu, Mark Whitaker, Anthony D Person, Vassili Kalabokis

## Abstract

Like sialylation, fucose usually locates at the non-reducing ends of various glycans on glycoproteins and constitutes important glycan epitopes. Detecting the substrate glycans of fucosyltransferases is important for understanding how these glycan epitopes are regulated in response to different growth conditions and external stimuli. Here we report the detection of these glycans via enzymatic incorporation of fluorescent tagged fucose using fucosyltransferases including FUT2, FUT6, FUT7, and FUT8 and FUT9. More specifically, we describe the detection of substrate glycans of FUT8 and FUT9 on therapeutic antibodies and the detection of high mannose glycans on glycoproteins by enzymatic conversion of high mannose glycans to the substrate glycans of FUT8. By establishing a series of precursor glycans, we also demonstrate the substrate specificities of FUT8. Furthermore, using simultaneous enzymatic incorporation of both fluorescent sialic acids and fluorescent fucoses, we demonstrate the interplay between fucosylation and sialylation.

## Introduction

Glycans are commonly found on cell membranes and secreted proteins. They are frequently terminated with sialic acids, negatively charged monosaccharides, and fucose, a deoxy hexosaccharide. For their unique physical properties, sialic acids and fucose are essential constituents of various glycan epitopes that are recognized by lectins and antibodies and involved in important biological roles ^1–4^.

Well-known fucosylated glycans include blood group H-antigen, Lewis X structures, and core fucosylated N-glycan. They are generated through various fucosyltransferases ^5^ (Fig. 1). H-antigen on red blood cells contains an α1-2 linked fucose introduced by FUT1 and FUT2 ^6^. Lewis X structure is a trisaccharide (Galβ1-4[Fucα1-3] GlcNAc) that has a fucose residue linked to a GlcNAc residue through an α1-3 linkage. Lewis X structure can be sialylated at the Gal residue to become sialyl-Lewis X structure (Neu5Acα2-3Galβ1-4[Fucα1-3] GlcNAc) that is the ligand for E-selectin and is essential for lymphocyte extravasation ^7^. The α-3 linked fucose on Lewis X and sialyl-Lewis X structures is introduced via several fucosyltransferases including FUT6, FUT7 and FUT9 ^8^. Among these enzymes, FUT7 is strictly active on sialyllactosamine ^9^, FUT9 is strictly active on lactosamine ^10^, and FUT6 is active on both structures ^11^. Fucosylation carried out by FUT9 is critical to ricin toxicity ^12^. Core-6 fucosylation on the innermost GlcNAc of N-glycan introduced by FUT8 ^13^ plays critical role in the antibody-dependent cellular cytotoxicity (ADCC) of therapeutic antibodies ^14^. For FUT8 substrate recognition, an unmodified β1-2 linked GlcNAc residue introduced to the α-3 arm of N-glycan by MGAT1 ^15^ is critical ^16^.

**Fig. 1.**
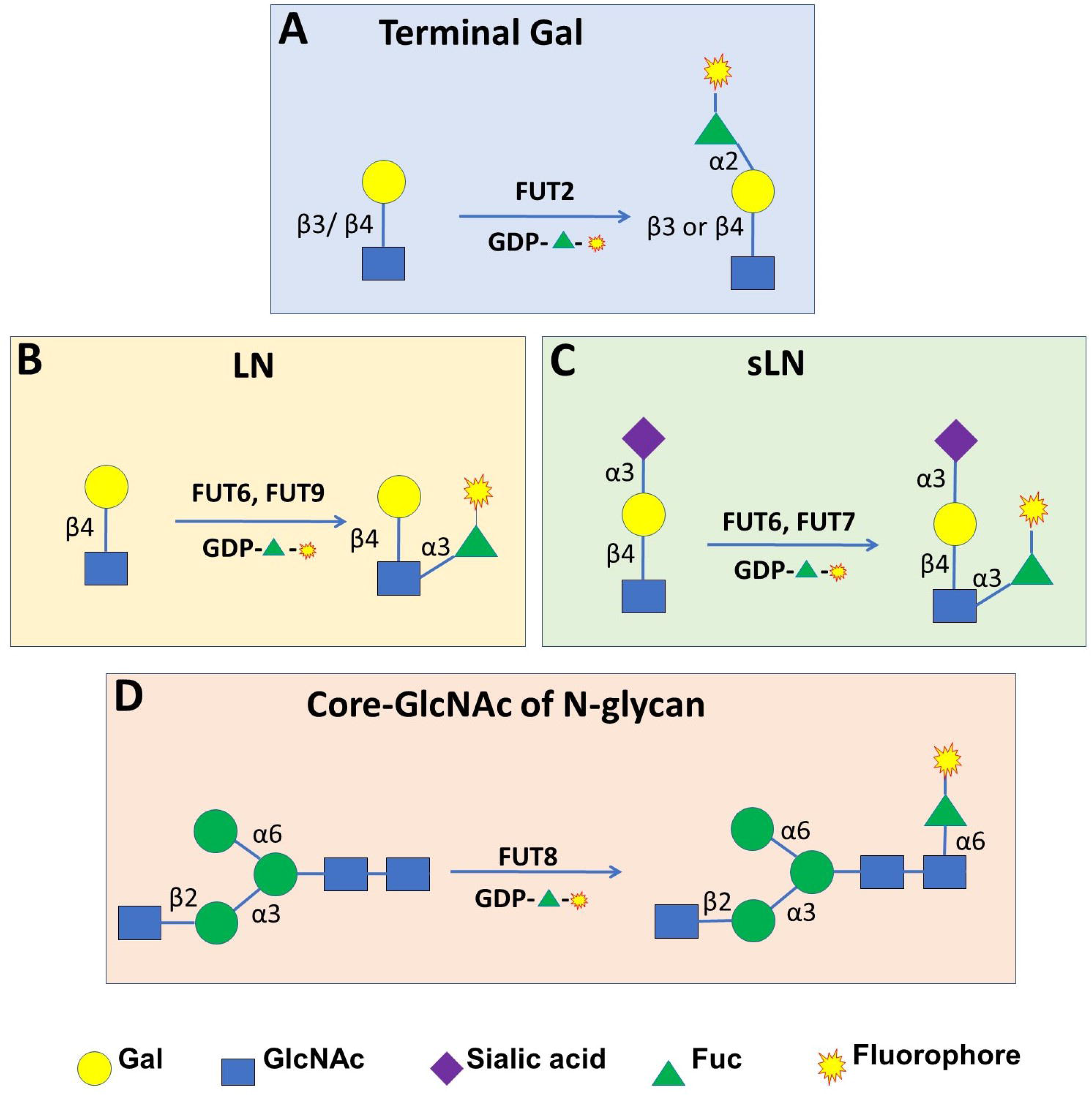
Substrate specificities of various fucosyltransferases and strategies for their substrate glycans labeling. **A**) FUT2 substrate specificity and its substrate glycan labeling. **B**) Strategy for labeling lactosamine (LN) by α-3 linkage specific FUT6 and FUT9. **C**) Strategy for labeling sialyl-lactosamine (sLN) by α-3 linkage specific FUT6 and FUT7. **D**) FUT8 substrate specificity and its substrate glycan labeling. FUT8 specifically introduces a fucose residue to the core-GlcNAc of an N-glycan that has a terminal β-2 linked GlcNAc on its α-3 arm.

For their important biological roles, cellular display of fucosylated glycan epitopes ^17^ must be tightly regulated ^4^. It is believed that this regulation is achieved via the establishment of precursor glycan pools and controlled expression of key FUTs. Upon environmental stimuli, cells can quickly convert the precursor glycans to functional epitopes via the action of these enzymes. Therefore, it is equally important to detect the glycan epitopes and their precursor glycans.

Previously, we described a direct fluorescent glycan labeling (DFGL) strategy to label and detect the substrate glycans of various sialyltransferases ^18^. In this report, we describe another DFGL strategy to label and detect the substrate glycans of some representative FUTs. The method was demonstrated on several well characterized glycoproteins, including fetal bovine fetuin^5^ that contains complex N-glycans and O-glycans, ribonuclease B^19^ that contains high mannose N-glycans, insect cell expressed recombinant H1N1 neuraminidase ^20^ that contains Man3 type high mannose N-glycan, and Cantuzumab ^21^, a therapeutic antibody that contains N-glycans. In addition, using enzymatic incorporation of fluorophore-tagged fucoses and sialic acids, we demonstrated dual labeling of N- and O-glycans on the cellular extracts of HEK293 cells and the interplay between FUT9 and N-glycan specific sialyltransferase ST6Gal1.

## Results

### Detection of Substrate Glycans of α-2 and α-3 Fucosyltransferases on Fetal Bovine Fetuin

To test whether we can detect the substrate glycans of α-2 and α-3 FUTs, we first prepared Cy5-conjugated GDP-Cy5-Fuc, and tested it as a donor substrate for FUT2, FUT6, FUT7 and FUT9 on fetal bovine fetuin and asialofetuin (Fig. 2A). The labeled samples were then separated on SDS-PAGE, followed by traditional protein gel imaging and fluorescent imaging. By comparing the images, it was found that these enzymes indeed can recognize Cy5-conjugated fucose (Cy5-Fuc) and label their substrate glycans. Specifically, we found that FUT2 and FUT9 can label asialofetuin, FUT7 can label fetuin and FUT6 can label both fetuin and asialofetuin. These results are consistent to the specificities of these enzymes reported in the literature ^8–11^.

**Fig. 2.**
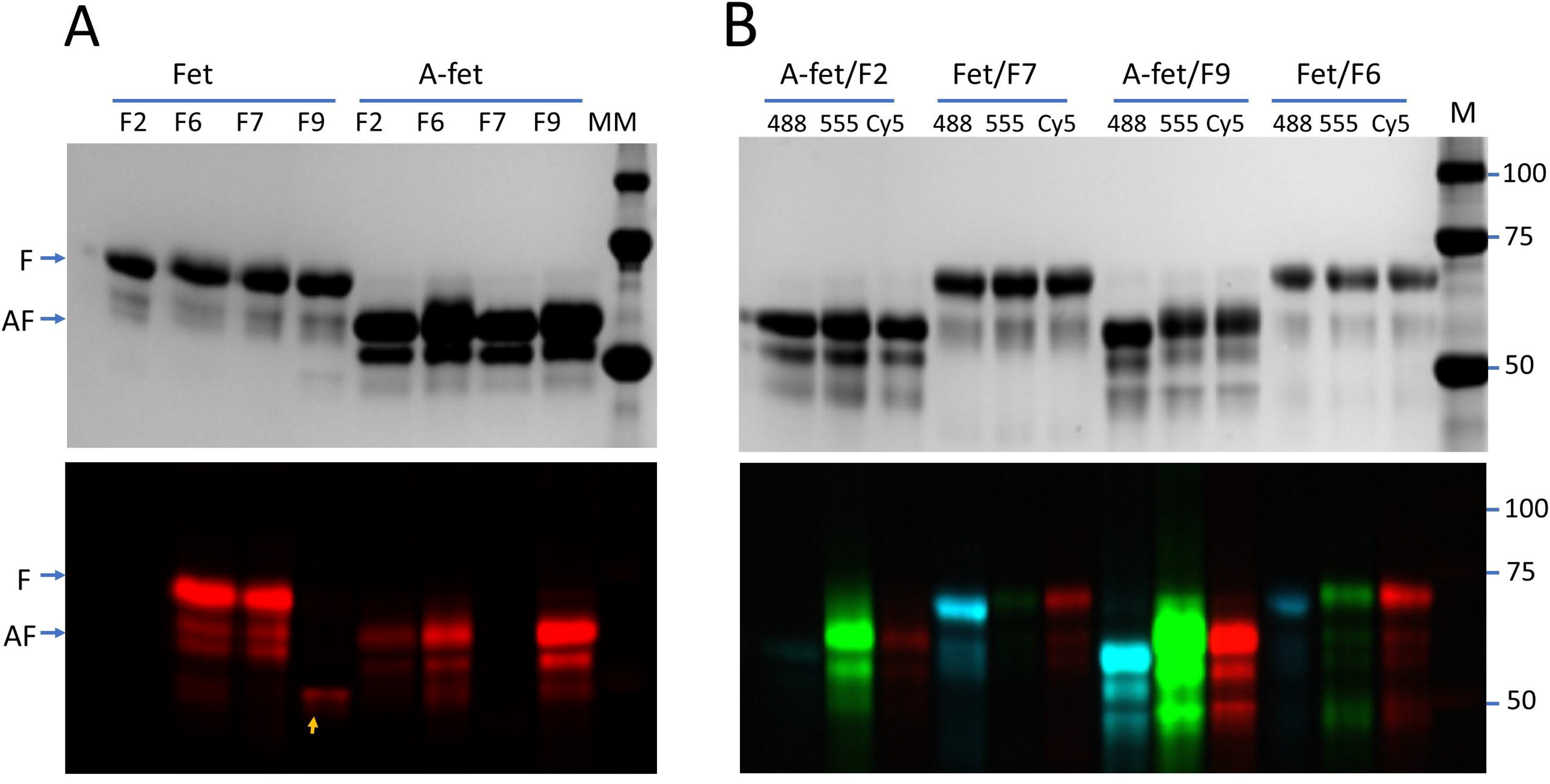
Probing substrate glycans of various fucosyltransferases on fetuin and asialofetuin. **A**) Probing substrate glycans of fucosyltransferases with Cy5. Samples of fetuin (Fet) or asialofetuin (A-fet) of fetal bovine origin were probed with FUT2(F2), FUT6(F6), FUT7(F7) and FUT9 (F9) in the presence of GDP-Cy5-fucose. FUT9 labeled itself in the gel (indicated by an arrow). **B**) Tolerance of Alexa-Fluor^®^555 (555), Alexa-Fluor^®^488 (488) and Cy5 by the indicated fucosyltransferases. All reactions were incubated at 37°C for 30 minutes and then separated on SDS-PAGE and imaged with silver staining (upper panels) and fluorescent imager (lower panels). M, molecular marker.

Next we tested the tolerance of the aforementioned FUTs for Cy5, AlexaFluor^®^ 488, and AlexaFluor^®^ 555 conjugated fucoses (Fig. 2B). We found that FUT2 preferred AlexaFluor^®^ 555, FUT7 preferred AlexaFluor^®^ 488, FUT9 preferred AlexaFluor^®^ 555, and, FUT6 showed no obvious preference among the three fluorescent fucoses (Table 1).

**Table 1.** Fucosyltransferase used in this study and their tolerance* for three fluorophores

| Fucosyltransferases | Acceptor glycan type | Alexa Fluor<sup>®</sup> 555 | Alexa Fluor<sup>®</sup> 488 | Cy5 |
| --- | --- | --- | --- | --- |
| FUT2 | Terminal Gal of lactosamine | +++ | + | + |
| FUT6 | GlcNAc of lactosamine, sialylation on terminal Gal is flexible | ++ | ++ | ++ |
| FUT7 | GlcNAc of sialylactosamine | + | ++ | + |
| FUT8 | Core GlcNAc of N-glycans | ++ | ++ | +++ |
| FUT9 | GlcNAc of lactosamine | +++ | +++ | +++ |

**Note:** \*The tolerance of a fluorophore-conjugated fucose as a donor substrate for a fucosyltransferase is based on the corresponding fluorescent intensity of the labeled fetuin or asialofetuin in Fig. 2, or labeled Cantuzumab in Fig. 3, and, is indicated by number of + sign. As the fluorescent intensity of each labeled band is also dependent on the abundance of the glycan acceptor on a target protein, comparison of the tolerance of different fluorophores should be limited to those under a same labeling enzyme.

### Probing the Status of Core-6 Fucosylation on Therapeutic Antibodies by FUT8

It is known that FUT8 can tolerate azido-fucose ^20^. Here we tested whether FUT8 can tolerate the three fluorescent fucoses and detect its substrate glycans on Cantuzumab that was prepared from FUT8 knockout cell line and the reference monoclonal antibody from the National Institute of Standards and Technology (NIST mAb, material 8671). NIST mAb is a humanized IgG1κ monoclonal antibody^22^. IgG antibodies are known to contain an N-glycan site on their heavy chains ^23^. Fig. 3A shows that significant amounts of Alexa-Fluor^®^ 555, Alexa-Fluor^®^ 488 and Cy5 conjugated fucoses were introduced into Cantuzumab but not to NIST mAb by FUT8. For comparison, the samples were also probed by FUT9, which showed consistent incorporation of the three dyes to both antibodies.

**Fig. 3.**
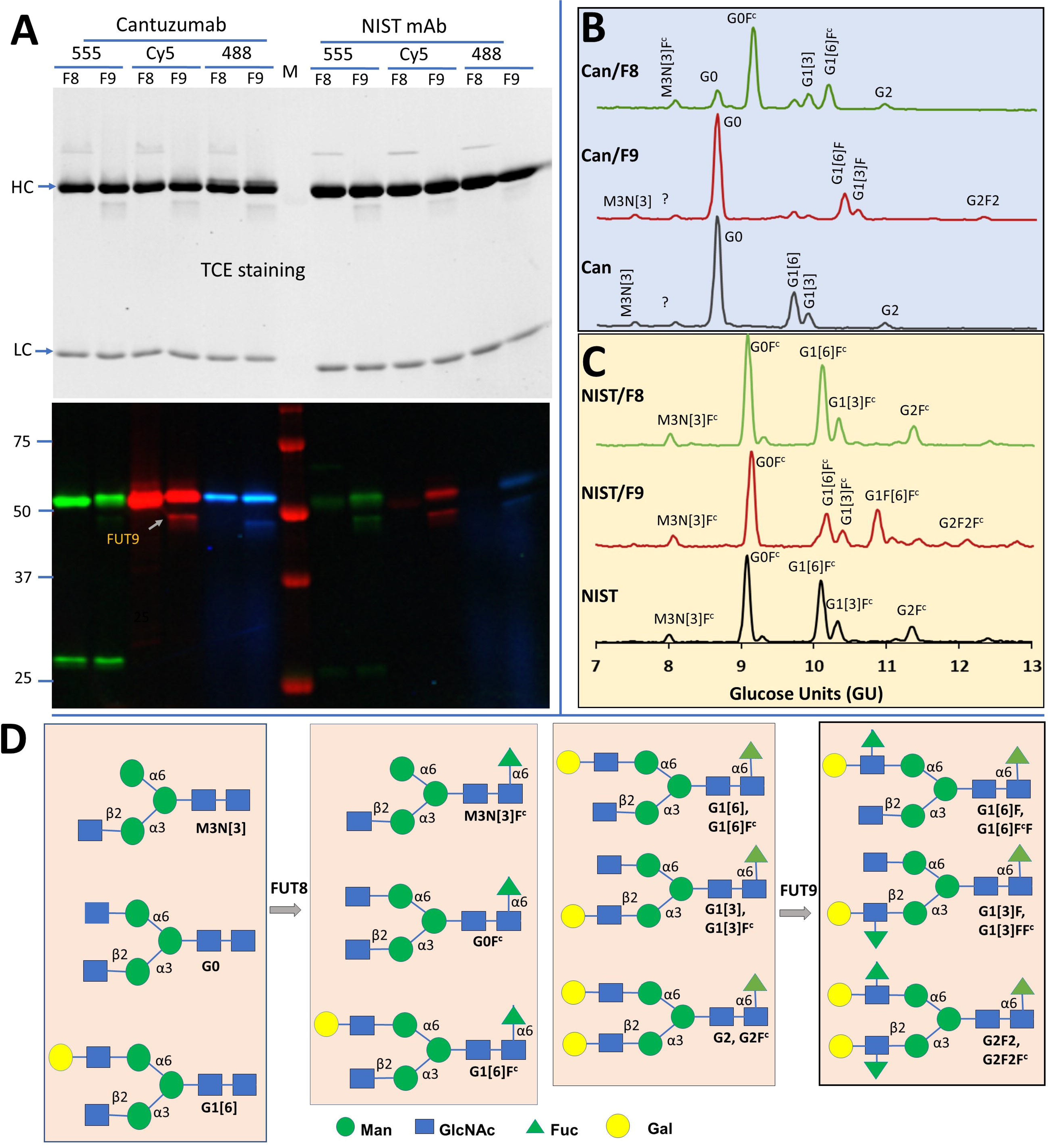
Probing core-6 fucosylation on antibodies by FUT8. **A**) Probing substrate glycans of FUT8 (F8) and FUT9 (F9) on Cantuzumab, an anti-Muc1 therapeutic antibody, and NIST reference mAb 8671 using GDP-Alexa-Fluor^®^555-fucose (555), GDP-Cy5-fucose (Cy5), and GDP-Alexa-Fluor^®^488-Fucose (488) as donor substrates. Cantuzumab was expressed in FUT8 knockout cells and is devoid of core-6 fucose. Molecular marker (MM) was a prestained Western molecular marker (BioRad). FUT9 (50 kDa) showed self-labeling. All reactions were separated on SDS-PAGE and imaged with trichloroethanol (TCE) staining (upper panel) and fluorescent imager (lower panel). Labeling on the light chains of the antibodies by Alexa-Fluor^®^555 is likely through non-specific staining due to the usage of unpurified GDP-Alexa-Fluor^®^555-fucose that contained free Alexa-Fluor^®^ 555. **B**) GlyQ analysis of the Cantuzumab samples prepared side-by-side for the samples in **A,** showing that FUT8 modified G0 and G1[6], and, FUT9 modified G1[6] and G1[3]. **C**) GlyQ analysis of the NIST mAb samples prepared side-by-side for the samples in **A,** showing that FUT8 had no modification on any glycan species and FUT9 modified G1[6]F^c^ and G2F^c^. **D**) The structures of the glycans analyzed in **B** and **C**.

To prove that the labeling was specific to respective enzyme substrate glycans, in a parallel experiment, glycans of *in vitro* fucosylated Cantuzumab and NIST mAb were analyzed on a Gly-Q™ Glycan Analysis System (Prozyme) (Fig. 3B). While FUT9 converted G1[6], G1[3] and G2 of Cantuzumab to G1[6]F, G1[3]F, and G2F2, respectively, FUT8 converted M3N[3], G0, and G1[6] of the antibody to M3N[3]F^c^, G0F^c^ and G1[6]F^c^, respectively, demonstrating the strict specificities of these two FUTs. Consistent to the labeling results of Fig. 3A, glycan analysis of NIST mAb suggested that G1[6]F^c^ and G2F^c^ were modified by FUT9 but no detectable substrate glycans for FUT8 were found (Fig. 3C). In fact, all peaks in the electropherogram of NIST mAb were found to be core-6 fucosylated. The structures of above mentioned glycans are shown in Fig. 3D.

### Detecting High Mannose Glycans by FUT8

High mannose N-glycans are related to serum clearance of therapeutic antibodies ^24^ and are frequently targeted in broad neutralizing antibody responses during human immunodeficiency viral infection ^25^. As such, detection of high-mannose glycans is particularly valuable. Here, we demonstrate a strategy to probe high mannose glycans using FUT8 and show the substrate specificity of FUT8. Bovine ribonuclease B (RNase B) is known to contain high-mannose glycans ^19^. To test whether we can detect high-mannose glycans on a glycoprotein, a sample of RNase B was first treated with α-1,3-mannosyl-glycoprotein 2-β-N-acetylglucosaminyltransferase (MGAT1) to introduce the α1-3-arm GlcNAc residue before labeling by FUT8 (Fig. 1D). The addition of α1-3-arm GlcNAc residue by MGAT1 resulted in strong labeling by FUT8, and further galactosylation and sialylation significantly reduced the labeling (left side of Fig. 4A). In addition, pretreatment of RNase B with unmodified fucose by FUT8 abolished the labeling completely, suggesting that the modification of fucose by Cy5 didn’t affect the substrate recognition by FUT8. As a positive control, an RNase B sample pretreated with MGAT1 and β-1,4-galactosyltransferase 1 (B4GalT1) was labeled with Cy5 conjugated sialic acid by ST6Gal1, which resulted in similar intensity of labeling by FUT8.

**Fig. 4.**
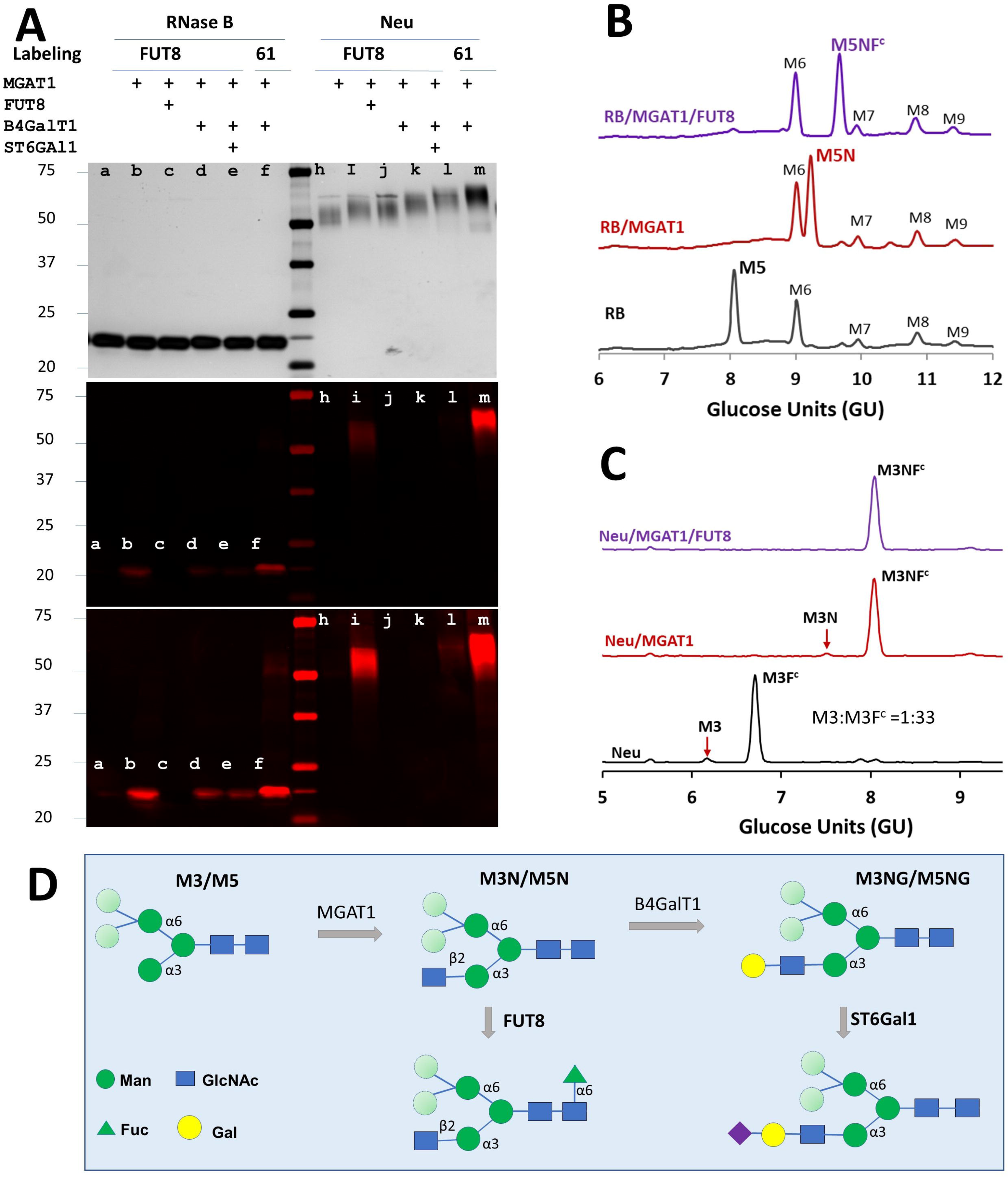
Detecting high mannose glycans on glycoproteins by FUT8. **A**) Detecting Man5 on RNase B and Man3 on recombinant H1N1 neuraminidase monomer (Neu). Samples were pretreated with MGAT1, FUT8, B4GalT1 and ST6Gal1 with their respective unmodified donor substrates (indicated with +). The pretreated samples were then labeled by FUT8 with GDP-Cy5-fucose or ST6Gal1 (61) with CMP-Cy5-Sialic acid. All samples were separated on SDS-PAGE and visualized by silver staining and fluorescent imaging. The middle and lower panels belong to a same image with different contrasts. Only the MGAT1 modified sample was strongly labeled by FUT8. **B**) Representative GlyQ analysis data of pretreated samples of RNase B (RB), showing that Man5 (M5) was sequentially modified by MGAT1 and FUT8. **C**) Representative GlyQ analysis data of pretreated samples of Neu, showing that both Man3 (M3) and M3F^c^ were modified by MGAT1, but only M3N was further modified by FUT8. **D**) Molecular structures of Man3 and Man5 and their derivatives for FUT8 and ST6Gal1 labeling. While the β2 linked GlcNAc introduced by MGAT1 on α3 arm is essential for FUT8 recognition, the mannose residues in lighter shade of green on α6 arm are flexible for substrate recognition. Further elongation of the α3 arm with B4GalT1 renders the glycan to be the substrate glycan for ST6Gal1. Only glycans involved in this experiment are depicted here.

To confirm that the glycans labeled by FUT8 and ST6Gal1 were high-mannose glycans, sequentially modified RNase B samples were analyzed with Gly-Q™ Glycan Analysis System. The results indicated that only Man5 (M5) was modified by FUT8 via MGAT1 (Fig. 4B) and ST6Gal1 via MGAT1 and B4GalT1 (Supplemental Fig.1). Since other high-mannose glycans including Man6, Man7, Man8, and Man9 can be converted to Man5 by α1,2 specific mannosidase ^26^ treatment, in theory, all these glycans can be detected by FUT8 as well.

To test whether Man3 glycan can be labeled, monomeric Sf21 cell expressed recombinant 1918 H1N1 influenza neuraminidase (Neu) that is known to contain both Man3 (M3) and core-6 fucosylated Man3 (M3F^c^) ^20^ was labeled by FUT8. Again, the sample was labeled significantly by FUT8 only after pretreatment with MGAT1 and the labeling was inhibited or abolished by additional pretreatment by B4GalT1 and ST6Gal1 (right side of Fig. 4A), again confirming that an unmodified GlcNAc introduced by MGAT1 on the α1,3-arm of high-mannose N-glycan is critical for FUT8 recognition. Meanwhile, since the difference between Man5 and Man3 is on their α1,6-arms, these results also proved that the α1,6-arms are flexible for FUT8 recognition. In contrast to labeling on RNase B, the signal of FUT8 labeled product was only a fraction of that of ST6Gal1 labeled product. To understand this difference and confirm that Man3 was indeed modified, sequentially modified Neu samples were subject to GlyQ analysis. It was found that the precursor substrate glycan for FUT8 (M3) on recombinant 1918 H1N1 influenza neuraminidase was only about 3% of that of the precursor substrate glycans for ST6Gal1 (both M3 and M3F^c^) (Fig. 4C), therefore explaining the difference on the signals labeled by FUT8 and ST6Gal1 in Fig. 4A. Similar results were obtained when the experiment was repeated on both monomeric and dimeric recombinant 1918 H1N1 influenza neuraminidase prepared in different batches (Supplemental Fig. 2).

### Interplay between Sialylation and Fucosylation Revealed by Simultaneous Labeling of Fucose and Sialic Acid

Since fucosylation and sialylation involve different donor substrates, it may be possible to label a common substrate glycan with fucosyltransferases and sialyltransferases simultaneously, thereby to study the interplay between these two families of enzymes. To test this idea, cytoplasmic extracts of HEK293 cells were labeled simultaneously by a sialyltransferase (ST6Gal1 or ST3Gal2) and a fucosyltransferase (FUT7 or FUT9) (Fig. 5). ST6Gal1 is active on terminal Galβ1-4GalNAc disaccharide on N-glycans ^27, 28^. ST3Gal2 is active on terminal Galβ1-3GalNAc disaccharide found on O-glycans ^28, 29^. To generate substrate glycans for FUT9, samples were pretreated with recombinant *C.p* neuraminidase to remove terminal sialic acids. The four enzymes exhibited distinctive labeling patterns, especially at bands around 10 kDa (likely to be labeled glycopeptides), demonstrating these enzymes recognize overlapping but distinct glycan substrates (Fig. 5B). More specifically, FUT9 and ST6Gal1 exhibited largely overlapping but not identical labeling patterns. When the extract was labeled simultaneously by FUT9 and ST6Gal1, the signal generated by either of the two enzymes was reduced compared to the signals generated by them individually (Fig. 5C), suggesting that the two enzymes have mutually exclusive relationship on their substrate recognition, *i.e.* sialylation by ST6Gal1 prevents fucosylation by FUT9 and vice versa. In contrast, ST3Gal2 showed no obvious interference to FUT9 labeling. ST6Gal1 and FUT7 exhibited very weak labeling on samples without neuraminidase treatment, suggesting that the proteins in the extracts were largely α2-6 sialylated and didn’t contain the substrate glycans for FUT7.

**Fig. 5.**
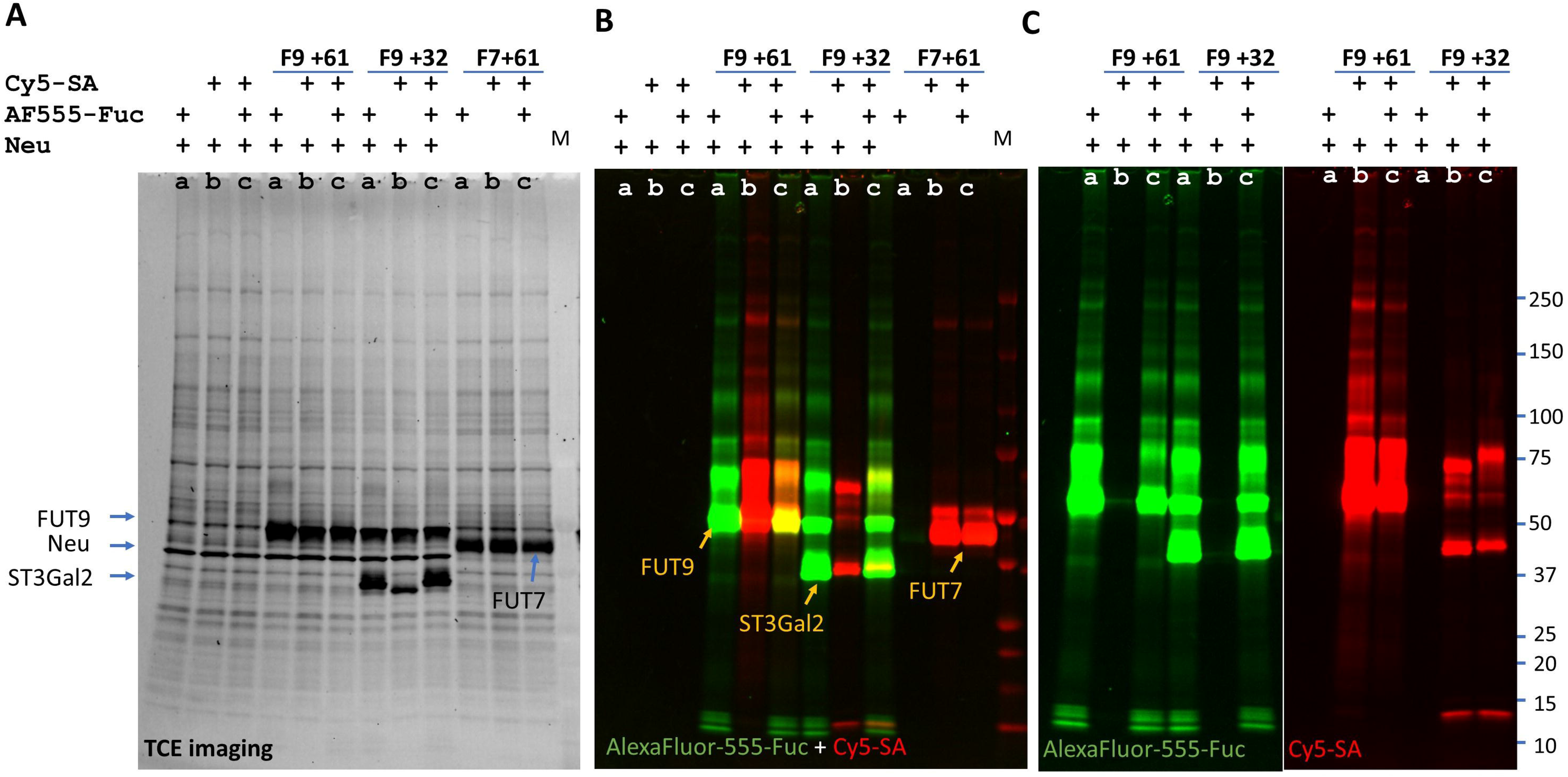
Simultaneous labeling by FUT9 (F9) or FUT7 (F7), and, ST6Gal1 (61) or ST3Gal2 (32) on HEK293 cell extracts. HEK293 cell extracts were labeled with Alexa-Fluor^®^ 555 conjugated fucose (AF555-Fuc) by an indicated fucosyltransferase (lane a) or Cy5 conjugated sialic acid (Cy5-SA) by an indicated sialyltransferase (lane b) or both (lane c). Recombinant *C.p* neuraminidase was added into some lanes as indicated to remove existing terminal sialic acids. All reactions were separated on SDS-PAGE. **A**) TCE image of the gel. **B**) Fluorescent image of the gel from both green and red channels. FUT9 labeled by itself, ST3Gal2 labeled by FUT9, and FUT7 labeled by ST6Gal1 are indicated by arrows. **C**) Single channel images of FUT9+ST6Gal1 and FUT9+ ST3Gal2 double labeling. M, prestained Protein Ladder (BioRad, visible from the red channel).

## Discussion

In this article, we demonstrated that Cy5, AlexaFluor^®^555 and AlexaFluor^®^ 488 conjugated fucoses are well tolerated by various FUTs. Using enzymatic incorporation of these fluorescent fucoses, we were able to reveal the presence of the substrate glycans of various FUTs on glycoproteins, particularly on therapeutic antibodies. We also demonstrated the detection of high mannose glycans on glycoproteins and the substrate specificities of FUT8. Furthermore, by enzymatic incorporation of fluorescent fucoses and fluorescent sialic acids simultaneously, we can study the interplay between fucosyltransferases and sialyltransferases.

Previously, we described direct fluorescent glycan labeling (DFGL) using enzymatic incorporation of fluorescent sialic acids ^18^. The current strategy is another version of DFGL that uses enzymatic incorporation of fluorescent fucoses. The two labeling strategies are similar in principle but have overlapping applications. In some cases, glycans can be labeled by either strategy. For example, terminal lactosamine can be labeled by either ST6Gal1 or FUT9, and high-mannose glycans can be converted to substrate glycans for either ST6Gal1 or FUT8 for labeling (Fig. 4). In some other cases, the substrate glycans can only be revealed by one strategy but not the other. For example, the status of core-6 fucosylation can only be revealed by FUT8 and the sialylation on core-1 O-glycan can only be revealed by ST3Gal1 or ST3Gal2. Labeling through fucosylation expands the range of glycans that can be labeled and provides further assurance when a substrate glycan can be labeled by both strategies.

Sialylation and fucosylation are common terminal modifications on various glycans. The interplay between these two modifications could determine some important biological properties of a cell. For examples, opposing actions of FUT9 and ST3Gal4 determines the sensitivity of a cell to the toxin ricin ^30^, and, the opposing actions of FUT2 and ST3Gals determines sialyl Lewis X expression ^31^. With these tools, we can study this kind of interplay rather conveniently. As an example, we demonstrated the mutually exclusive relationship between the sialylation by ST6Gal1 and the fucosylation by FUT9. Since sialylation by ST6Gal1 creates the receptors for H1N1 influenza virus^32^, the counteractive action by FUT9 could mitigate the susceptibility of a cell to the virus. In contrast, no obvious interplay was found between ST3Gal2 and FUT9, which is not surprising as these two enzymes recognize different substrate glycans.

Since the fucosyltransferases investigated in this report can tolerate Cy5 and AlexaFluor fluorophores that are very different on structures, it might be possible to use these enzymes to conjugate other types of functional groups such as drugs to glycoproteins on cell surface as a way for drug targeting.

## Material and methods

Recombinant FUT2, FUT6, FUT8, FUT9, MGAT1, B4GalT1, ST6Gal1, H1N1 viral neuraminidase, *C. perfringens* Neuraminidase and GDP-Azido-Fucose were obtained from Bio-Techne. Cantuzumab, an anti-Muc1 therapeutic antibody, was purchased from Creative Biolabs. NIST monoclonal antibody reference material 8671 was purchased directly from the National Institute of Standards and Technology. Alkyne-Alexa Fluor^®^ 488, alkyne-Alexa Fluor^®^ 555 were purchased from Thermo Fisher Scientific. Cy5-alkyne, RNase B, fetal bovine fetuin and asialofetuin and all other chemical reagents were purchased from Sigma-Aldrich.

### Preparation of fluorescent conjugated GDP-fucose

Fluorophore conjugated GDP-fucoses were prepared by incubating equivalent GDP-Azido-Fucose (CMP-N_3_-Fuc) and alkyne-conjugated fluorophores via copper (I)-catalyzed azide-alkyne cycloaddition. For example, 5 mM of GDP-N_3_-Fuc was mixed with 5 mM of Cy5-alkyne in the presence of 0.1 mM of Cu^2+^ and 1 mM of ascorbic acid. The reaction was kept at room temperature for 2 hours. The fluorophore conjugated GDP-fucoses were purified on a HiTrap^®^ Q HP (GE Healthcare) column and eluted with a 0-100% gradient of NaCl elution buffer (300 NaCl, 25 mM Tris at pH 7.5). The fluorophore conjugated GDP-fucoses were collected based on color exhibition and UV absorption as the conjugated GDP-fucoses were vivid in color and had UV absorption at 260 nm. Alexa Fluor^®^ 555 conjugated GDP-fucose, Alexa Fluor^®^ 488 conjugated GDP-fucose, Cy5 conjugated GDP-fucose were prepared and purified and finally concentrated to > 0.1 mM by a speed-vacuum concentrator.

### Fluorescent labeling of glycoproteins using fucosyltransferases

For a typical labeling reaction, 1 to 5 μg of a target protein was mixed with 0.2 nmol fluorophore conjugated GDP-fucose and 0.2 μg of a fucosyltransferase in 30 μL 25 mM Tris pH 7.5, 10 mM MnCl_2_. The mixture was incubated at 37°C for 30 minutes. The reaction was then separated by sodium dodecyl sulfate–polyacrylamide gel electrophoresis (SDS-PAGE) and the gel was directly imaged using a fluorescent imager FluorChem M (ProteinSimple, Bio-techne) and followed by imaging with traditional protein imaging methods such as silver staining or trichloroethanol (TCE) staining.

### GlyQ analysis

All samples for GlyQ analysis were prepared analyzed according to the manufacture’s protocol.

