## Supplemental Fig. 1 and 2 for "Detecting substrate glycans of fucosyltransferases on glycoproteins with fluorescent fucose"

### Slide 1
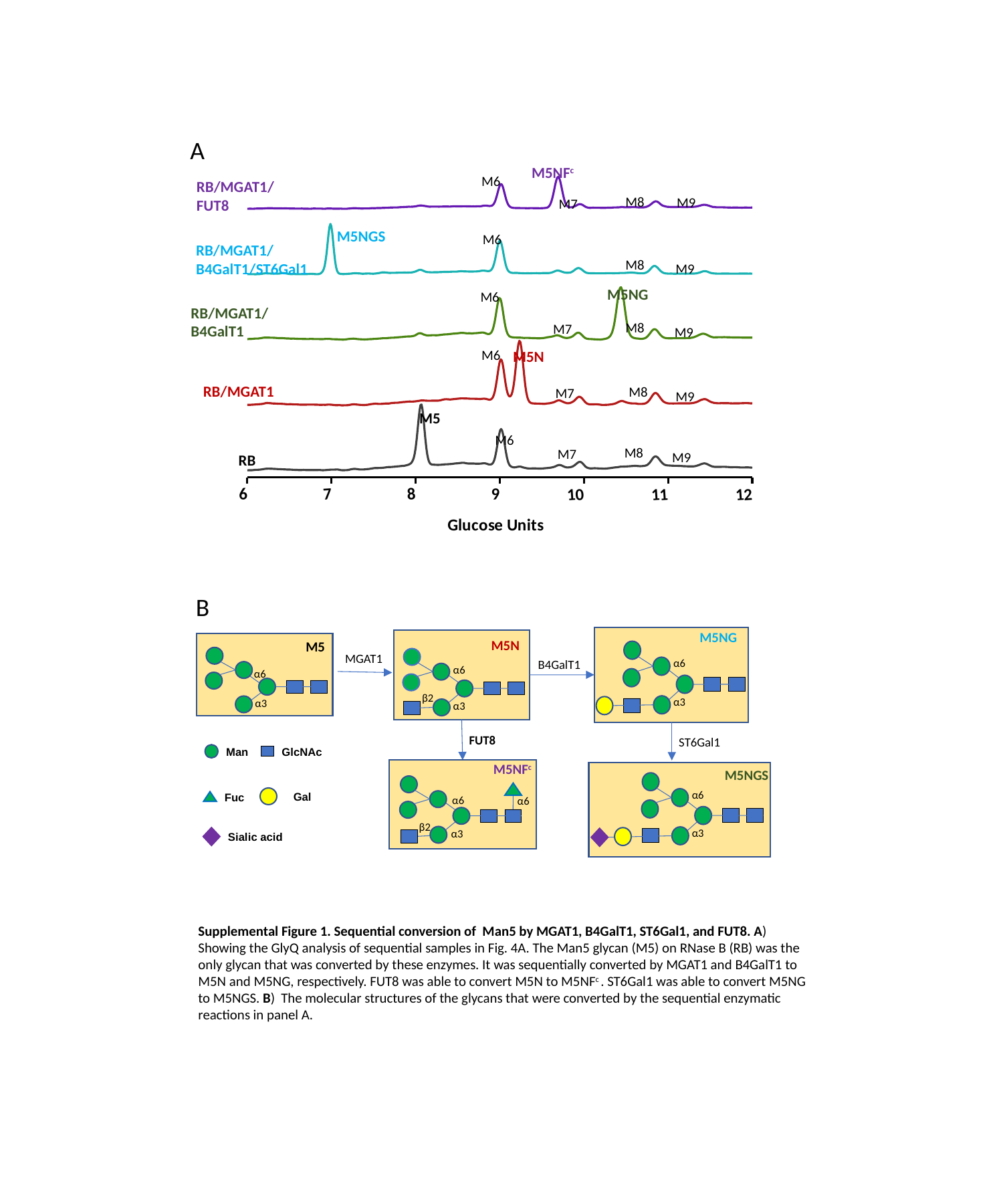

A
M5NF^c^
#### Chart
| Category | | | | | |
|---|---|---|---|---|---|M6
RB/MGAT1/
FUT8
M8
M9
M7
M5NGS
M6
RB/MGAT1/
B4GalT1/ST6Gal1
M8
M9
M5NG
M6
RB/MGAT1/
B4GalT1
M8
M7
M9
M6
M5N
RB/MGAT1
M8
M7
M9
M5
M6
M8
M7
M9
RB
B
M5NG
M5N
M5
α6
α3
MGAT1
α6
β2
α3
B4GalT1
α6
α3
FUT8
ST6Gal1
Man
GlcNAc
M5NF^c^
α6
α6
β2
α3
M5NGS
α6
α3
Gal
Fuc
Sialic acid
Supplemental Figure 1. Sequential conversion of Man5 by MGAT1, B4GalT1, ST6Gal1, and FUT8. A) Showing the GlyQ analysis of sequential samples in Fig. 4A. The Man5 glycan (M5) on RNase B (RB) was the only glycan that was converted by these enzymes. It was sequentially converted by MGAT1 and B4GalT1 to M5N and M5NG, respectively. FUT8 was able to convert M5N to M5NF^c^ . ST6Gal1 was able to convert M5NG to M5NGS. B) The molecular structures of the glycans that were converted by the sequential enzymatic reactions in panel A.

### Slide 2
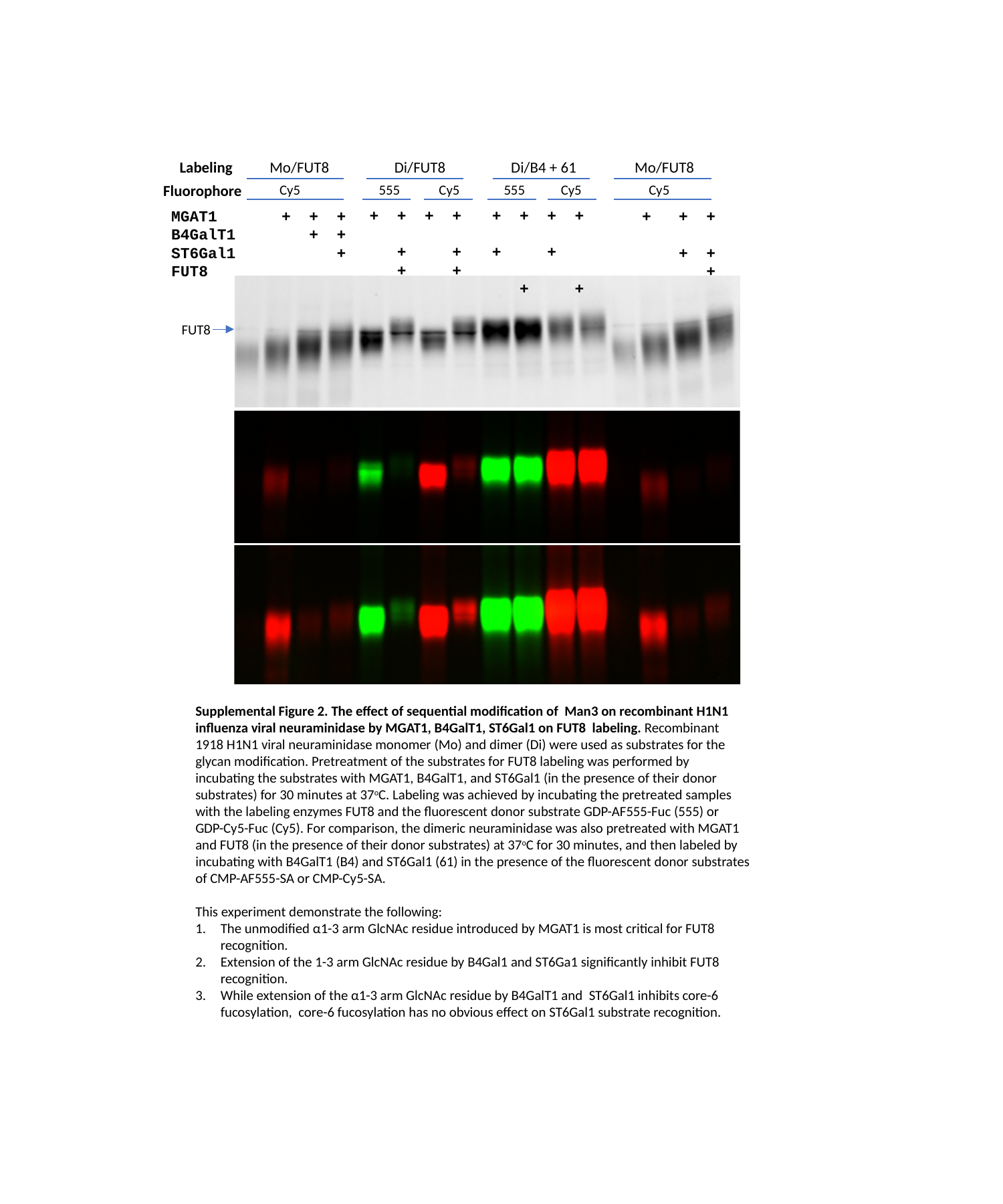

Labeling
 Mo/FUT8
Di/FUT8
Di/B4 + 61
Mo/FUT8
Cy5
555
Cy5
555
Cy5
Cy5
Fluorophore
+ + + +
 + +
 + +
+ + + +
+ +
 + +
 + + +
 + +
 +
MGAT1 + + +
B4GalT1 + +
ST6Gal1 +
FUT8
FUT8
Supplemental Figure 2. The effect of sequential modification of Man3 on recombinant H1N1 influenza viral neuraminidase by MGAT1, B4GalT1, ST6Gal1 on FUT8 labeling. Recombinant 1918 H1N1 viral neuraminidase monomer (Mo) and dimer (Di) were used as substrates for the glycan modification. Pretreatment of the substrates for FUT8 labeling was performed by incubating the substrates with MGAT1, B4GalT1, and ST6Gal1 (in the presence of their donor substrates) for 30 minutes at 37^o^C. Labeling was achieved by incubating the pretreated samples with the labeling enzymes FUT8 and the fluorescent donor substrate GDP-AF555-Fuc (555) or GDP-Cy5-Fuc (Cy5). For comparison, the dimeric neuraminidase was also pretreated with MGAT1 and FUT8 (in the presence of their donor substrates) at 37^o^C for 30 minutes, and then labeled by incubating with B4GalT1 (B4) and ST6Gal1 (61) in the presence of the fluorescent donor substrates of CMP-AF555-SA or CMP-Cy5-SA.
This experiment demonstrate the following:
The unmodified α1-3 arm GlcNAc residue introduced by MGAT1 is most critical for FUT8 recognition.
Extension of the 1-3 arm GlcNAc residue by B4Gal1 and ST6Ga1 significantly inhibit FUT8 recognition.
While extension of the α1-3 arm GlcNAc residue by B4GalT1 and ST6Gal1 inhibits core-6 fucosylation, core-6 fucosylation has no obvious effect on ST6Gal1 substrate recognition.
